# Homeostasis in Networks with Multiple Inputs

**DOI:** 10.1101/2022.12.07.519500

**Authors:** João Luiz de Oliveira Madeira, Fernando Antoneli

## Abstract

Homeostasis, also known as adaptation, refers to the ability of a system to counteract persistent external disturbances and tightly control the output of a key observable. Existing studies on homeostasis in network dynamics have mainly focused on ‘perfect adaptation’ in deterministic single-input single-output networks where the disturbances are scalar and affect the network dynamics via a pre-specified input node. In this paper we provide a full classification of all possible network topologies capable of generating infinitesimal homeostasis in arbitrarily large and complex multiple-input parameter networks. Working in the framework of ‘infinitesimal homeostasis’ allows us to make no assumption about how the components are interconnected and the functional form of the associated differential equations, apart from being compatible with the network architecture. Remarkably, we show that there are just three distinct ‘mechanisms’ that generate infinitesimal homeostasis. Each of these three mechanisms generates a rich class of well-defined network topologies – called *homeostasis subnetworks*. Most importantly, we show that these classes of homeostasis subnetworks provides a topological basis for the classification of ‘homeostasis types’: the full set of all possible multiple-input parameter networks can be uniquely decomposed into these special homeostasis subnetworks. We build on previous work that treated the cases of single-input node and multiple-input node, both with a single scalar input parameter. Furthermore, we identify a new phenomenon that occurs in the multiparameter setting, that we call *homeostasis mode interaction*, in analogy with the well-known characteristic of multiparameter bifurcation theory.

## 1 Introduction

A homeostatic process is characterized by the following property: approximately zero steady-state error to external disturbance, which means that an observable of interest is tightly controlled. Homeostasis is biologically important because it protects organisms against changes induced by the environment. A familiar example is thermoregulation, where the body temperature of an organism remains roughly constant despite variations in its environment [40]. Another example is a biochemical reaction network, where the equilibrium concentration of some important biochemical molecule might not change much when the substrate concentration changes [48]. Further examples include regulation of cell number and size [36], sleep control [56], and expression level regulation in housekeeping genes [2].

Homeostasis can be mathematically defined as follows (see Section 2.1). Consider a dynamical system depending on an input parameter ℐ which varies over an open interval]ℐ_1_, ℐ_2_[of external stimuli. Suppose there is a family of equilibrium points *X*(ℐ) and an observable *ϕ* such that the *input-output function z*(ℐ) = *ϕ*(*X*(ℐ)) is well-defined on]ℐ_1_, ℐ_2_[. In this situation, we say that the system exhibits *homeostasis* if, under variation of the input parameter ℐ, the input-output function *z*(ℐ) remains ‘approximately constant’ over the interval of external stimuli.

There are two formulations of ‘approximately constant’ often considered by researchers. The first one, called *perfect homeostasis*, where the input-output function is required to be constant over the interval of external stimuli. This strict version of homeostasis is widely studied in control engineering and synthetic biology under the name ‘perfect adaptation’ (cf. [1,4,18,32,37,39]). *Perfect adaptation* is defined as the ability of a system to reset to its pre-stimulated output level, called *set point*, after responding to an external disturbance. It is obvious that this condition is equivalent to the requirement that the input-output function is constant. The more general condition of *near-perfect homeostasis*, where the input-output function is required to be within a ‘narrow’ range when the input parameter varies over the interval of external stimuli. This notion of homeostasis has appeared under the names *near-perfect adaptation* [1, 17, 39] and *imperfect adaptation* [6, 7]. Despite some contribution in this direction [8], it has remained relatively unexplored.

The notion of perfect homeostasis is a stringent requirement appropriate for the design of engineering systems but is biologically unmotivated. In biology, homeostasis is an emergent property of the system, not a preset target value, and the input-output function is only approximately constant. On the other hand, near perfect homeostasis is a very flexible qualitative property, that must be made quantitative by introducing an artificial parameter to control the range of the variation of the output.

Golubitsky and Stewart [21] proposed an intermediate notion of homeostasis: a system exhibits *infinitesimal homeostasis* if 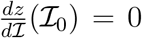 for some input value ℐ_0_ ∈]ℐ_1_, ℐ_2_ [. The vanishing of the derivative of *z* at ℐ_0_ implies that ℐ_0_ is a *critical point* of the input-output function and hence *z* is ‘approximately constant’ near ℐ_0_. Moreover, it is possible to find quantitative estimates on the size of the interval]ℐ_1_, ℐ_2_[where *z*(ℐ) stays within *z*(ℐ_0_) ± *δ*, for a given *δ >* 0 (see [23] for details). As we shall see in a moment, there are some additional advantages in adopting this point of view, besides providing a plausible notion of near-perfect homeostasis.

Nijhout, Reed, and Best [5, 41–46] among others, have shown that homeostasis is an important phenomenon in biochemical reaction networks. In a biochemical network, each node represents the concentration of a chemical substrate and each arrow denotes a chemical interaction between the molecules at the head and tail of the arrow. In an input-output network formulation, one node is designated as the input node *ι* and another is designated as the output node *o*. The modeling assumes that some external stimuli (represented by a parameter ℐ) affects the network dynamics only at the input node, and the end result of computation by the network dynamics is the value of the output node. In this setting, there is a canonical choice for observable *ϕ*: it is the coordinate function of the output node.

Motivated by these examples, Wang et al. [55] introduced the notion of ‘abstract input–output network’ and devised a scheme for the classification of ‘infinitesimal homeostasis types’ in such networks. In a precise way, the homeostasis types of a given network correspond to the possible ‘mechanisms’ that are responsible for the occurrence of infinitesimal homeostasis in that network. The results of [55] apply to the case of single-input single-output networks where the external stimuli can only affect the input node via a single input parameter.

Even though single-input single-output networks are quite popular in many engineering domains [1,4,7,37], the single input (and single input parameter) assumption seems unrealistic in biology, as disturbances that arise are typically very complex and do not have a single well-defined entry point. Maderia and Antoneli [38] extended the classification of [55] to the setting of multiple input nodes affected by a single input parameter. Using this extended theory they were able to completely work out the homeostasis types of a representative model for bacterial chemotaxis [11, 54].

It is not difficult to conceive that homeostasis can occur on variation of two or more input parameters [27]. An important biological example is homeostasis of extracellular dopamine (eDA) in response to variation in the activities of the enzyme tyrosine hydroxylase (TH) and the dopamine transporters (DAT) [5]. The full generalization of the infinitesimal homeostasis formalism of [21] to the multiple input parameter case was carried out by Golubitsky and Stewart [22].

In the setting of perfect homeostasis, Tang et al. [50], already considered the case of multiple input parameters and obtained the multidimensional algebraic condition for homeostasis under the name ‘cofactor condition’. The single input version of this algebraic condition has been obtained by several authors: [37] for three-node networks, [4] under the name ‘RPA equation’. See [21,24,38,48,55] for the treatment in the context of infinitesimal homeostasis (see Section 2.2).

### 1.1 Homeostasis Types in Multiparameter Networks

In this paper we further build on the theory of [38,55] to completely solve the problem of classifying the homeostasis types for input-output networks with multiple input nodes and multiple input parameters (Sec. 2.2). More precisely, given an input-output network with multiple input parameters we show that one can consider each parameter at a time. Thus, effectively reducing the problem of classification of homeostasis types to the single input parameter case (still with multiple input nodes), that has been completely solved in [38]. Afterwards, we show how to combine these partial classifications for the single input parameter cases into an algorithm that provides the full classification on the multiple input parameter setting (Sec. 2.7).

The main discovery of [38, 55] is that, in a given abstract input-output network, there is a finite number of ‘distinct mechanisms’ that may cause homeostasis, i.e., may force the derivative of the input-output function to vanish (at a fixed value of the input parameter). These ‘distinct mechanisms’, called ‘homeostasis types’, bijectively correspond to a specific subnetworks of the abstract input-output network. These subnetworks, called *homeostasis subnetworks* can be characterized in purely topological terms. Hence, one can write down an algorithm that completely lists all the homeostasis subnetworks of any abstract input-output network.

In the single-input single-output theory of [55] the homeostasis subnetworks can be divided into two classes: *structural* and *appendage*. The structural subnetworks correspond to feedforward mechanisms and the appendage subnetworks correspond to feedback mechanisms. These are called, respectively, ‘opposer modules’ and ‘balancer modules’ in the control theoretic literature [4, 7].

In [38] the single-input single-output theory is generalized, for the first time, to account for multiple input nodes. The main result is that everything from the single-input case generalizes, except that there is a new class of homeostasis subneworks, called *input counterweight* : every abstract multiple input node network has exactly one input counterweight subnetwork, which can be topologically characterized, as well. Hence, one can extend the algorithm for listing all the homeostasis subnetworks to the case of multiple input nodes.

The final generalization obtained here, allows one to completely classify homeostasis subnetworks of multiple input parameters ‘core’ network (Secs. 2.3, 3.1). In the multiparameter setting two new phenomena arise. The first phenomenon is related to being able to consider one parameter at a time. More specifically, given a scalar input parameter ℐ_*M*_ we define a unique subnetwork, called ℐ_*M*_ *-specialized subnetwork*, that contains all the homeostasis subnetworks associated to the input parameter ℐ_*M*_ (Sec. 2.4). Then we can divide the homeostasis subnetworks into two cases: (i) a *pleiotropic subnetwork* is contained in all ℐ_*M*_ - specialized subnetworks, (ii) a *coincidental subnetwork* is not contained in all ℐ_*M*_ - specialized subnetworks. Then we show that the class of a pleiotropic subnetwork can only be structural or appendage (Secs. 2.5, 3.2). Whereas the class of a coincidental subnetwork can be structural, appendage or input counterweight (Sec. 2.6).

The second new phenomenon of multiparameter setting is related to the occurrence of overlapping between coincidental subnetworks contained in distinct ℐ_*M*_ - specialized subnetworks. These non-trivial interactions between homeostasis subnetworks in multiparameter networks leads to the notion of *homeostasis mode interaction* or *higher codimension homeostasis*. The notion of mode interaction is familiar in bifurcation theory. In a steady-state bifurcation the eigenvectors of the linearized equation corresponding to simple eigenvalues are called *modes*. A mode whose eigenvalues lie on the imaginary axis is said to be *critical*. Generically, it is expected that a one-parameter system has only one critical mode. However, in systems with more than one parameter, one expects multiple critical modes. The secondary bifurcation that may arise in nonlinear systems near a parameter value at which there are multiple critical modes are thought of as resulting from a nonlinear interaction of several critical modes. This process is called *mode interaction* and the parameter values at which there are multiple critical modes are called *higher codimension bifurcation points*. We assert that there is an analogue process in the context of infinitesimal homeostasis (Ex. 2.12, 2.13, 2.14). Duncan et al. [13] investigates the appearance of codimension-two homeostasis mode interaction in a slightly different setting. Nevertheless, [13] already found a rich scenario with higher degeneracy of the homeostasis point or the occurrence of a steady-state bifurcation at the same time.

We end this section by briefly mentioning other aspects of the infinitesimal homeostasis approach that we will not pursue further: general network dynamical systems formalism (network admissible systems) [9, 19, 23], singularity theory network-preserving changes of coordinates [3, 20–22], biological robustness [29, 31, 33–35, 47, 51–53], numerical discovery and continuation [12, 16, 26], classification of infinitesimal homeostasis in small ‘core networks’ [28], infinitesimal homeostasis for limit cycles [57] and miscellaneous applications to biology [2, 24, 25, 38].

## 2 Homeostasis in Multiparameter Networks

In this section we give all essential definitions and state the main results of the paper.

### 2.1 Dynamical Theory of Infinitesimal Homeostasis

Golubitsky and Stewart proposed a mathematical method for the study of homeostasis based on dynamical systems theory [21, 22] (see the review [24]). In this framework, one consider a system of differential equations

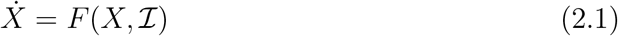

where *X* = (*x*_1_,, *x*_*n*_) ∈ ℝ^*n*^ is the vector of state variables and ℐ = (ℐ_1_, …, ℐ_*N*_) ∈ ℝ^*N*^ is the vector of external input parameter.

Suppose that (*X*\*, ℐ*) is a linearly stable equilibrium of (2.1). By the implicit function theorem, there is a function 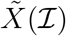 defined in a neighborhood of ℐ* such that 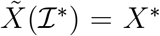 and 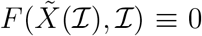. See [30] for results on the generic existence and robustness of 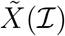.

A smooth function *ϕ* : ℝ^*n*^ × ℝ^*N*^ → ℝ is called an *observable*. Define the *input-output function z* : ℝ^*N*^ → ℝ associated to *ϕ* and 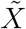 as 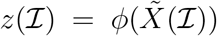. The input-output function allows one to formulate several definitions that capture the notion of homeostasis (see [1, 21, 22, 37, 50]).

#### Definition 2.1.

Let *z*(ℐ) be the input-output function associated to a system of differential equations (2.1). We say that *z*(ℐ) exhibits

a. *Perfect Homeostasis* on an open set Ω ⊆ ℝ^*N*^ if

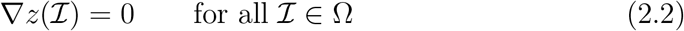 That is, *z* is constant on Ω.
b. *Near-perfect Homeostasis* relative to a *set point* ℐ_s_ on an open set Ω ⊆ ℝ^*N*^ if, for a fixed *δ*,

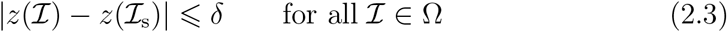 That is, *z* stays within the range *z*(ℐ_s_) ± *δ* over Ω.
c. *Infinitesimal Homeostasis* at the point ℐ_c_ on an open set Ω ⊆ ℝ^*N*^ if

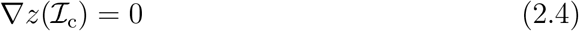 That is, ℐ_c_ is a *critical point* of *z*. ◊

It is clear that perfect homeostasis implies near-perfect homeostasis, but the converse does not hold. Inspired by Nijhout, Reed, Best et al. [5, 45], Golubitsky and Stewart [21, 22] introduced the notion of infinitesimal homeostasis that is intermediate between perfect and near-perfect homeostasis. It is obvious that perfect homeostasis implies infinitesimal homeostasis. On the other hand, it follows from Taylor’s theorem that infinitesimal homeostasis implies near-perfect homeostasis in a neighborhood of ℐ_c_ [23]. It is easy to see that the converse to both implications is not generally valid (see [48]).

The notion of infinitesimal homeostasis allows one to apply the tools from singularity theory. For instance, by considering higher degeneracy conditions, in addition to (2.4), one is lead distinct forms of infinitesimal homeostasis that can be classified by elementary catastrophe theory (see [21, 22] for details). Finally, when combined with coupled systems theory [19] the formalism of [21, 22, 24] becomes very effective in the analysis of model equations.

### 2.2 Multiparameter Input-Output Networks

A *multiple input parameter input-output network*, or simply *multiparameter input-output network*, is a network 𝒢 with *n* distinguished *input nodes ι* = {*ι*_1_, *ι*_2_, …, *ι*_*n*_}, all of them associated to at least one input parameter ℐ_*M*_, *M* = 1, …, *N*, one distinguished *output node o*, and *r regulatory nodes ρ* = {*ρ*_1_, …, *ρ*_*R*_}. The associated network systems of differential equations have the form

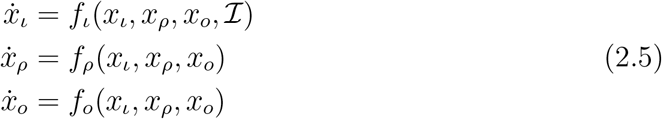

where ℐ = (ℐ_1_, …, ℐ_*N*_) ∈ ℝ^*N*^ is an *external multidimensional input parameter* and *X* = (*x*_*ι*_, *x*_*ρ*_, *x*_*o*_) ∈ ℝ^*n*^ × ℝ^*r*^ × ℝ is the vector of state variables associated to the network nodes.

We write a vector field associated with the system (2.5) as

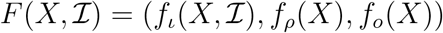

and call it an *admissible vector filed* for the network 𝒢.

Let 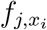 denote the partial derivative of the *j*^*th*^ node function *f*_*j*_ with respect to the *i*^*th*^ node variable *x*_*i*_. We make the following assumptions about the vector field *F* throughout:

a. The vector field *F* is smooth and has a linearly stable equilibrium at (*X*\*, ℐ*). Therefore, by the implicit function theorem, there is a function 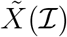 defined in a neighborhood of ℐ* such that 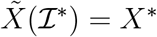 and 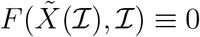.
b. The partial derivative 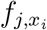 can be non-zero only if the network 𝒢 has an arrow *i* → *j*, otherwise 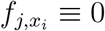.
c. Only the input node coordinate functions 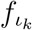 depend on at least one of the coordinates of the external input parameter ℐ and the partial derivative of 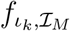 generically satisfies

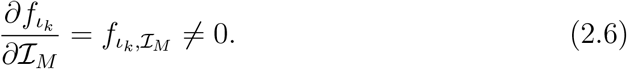

for some *M* = 1, …, *N*.

The network structure provides a distinguished class of observables for an admissible system, namely, the state variables. In the particular case of an input-output network the observable of interest is given by the output state variable *x*_*o*_.

#### Definition 2.2.

Let 𝒢 be a multiparameter input-output n(etwork and *F* be a)family admissible vector fields with an equilibrium point 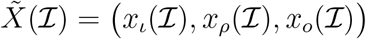. The mapping ℐ ⟼ *x*_*o*_(ℐ) is called the *input-output function* of the network 𝒢, relative to the family of equilibria 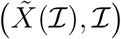.

Infinitesimal homeostasis in a multiparameter input-output network is given by the critical points of *x*_*o*_(ℐ), namely, the zeros of the gradient vector

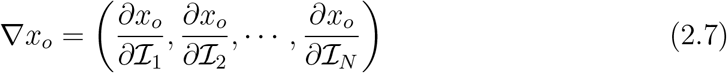

By a straightforward application of Cramer’s rule Wang et al. [55] obtained a determinant formula for the derivative of the input-output function in the single-input single-output case. Madeira and Antoneli [38] generalized the determinant formula of [55] to the case of multiple input nodes networks. In the following we further generalize the determinant formula of [38] to the case of multiple input parameters networks.

Let *J* be the (*n* + *r* + 1) × (*n* + *r* + 1) Jacobian matrix of an admissible vector field *F* = (*f*_*ι*_, *f*_*σ*_, *f*_*o*_), that is,

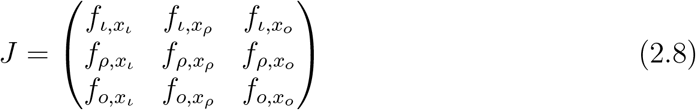

For each 1 ≤ *M* ≤ *N*, consider the (*n*+*r* +1)×(*n*+*r* +1) matrix ⟨ *H*_*M*_ ⟩ obtained from *J* by replacing the last column by 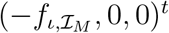, is called ℐ_*M*_ *-generalized homeostasis matrix* :

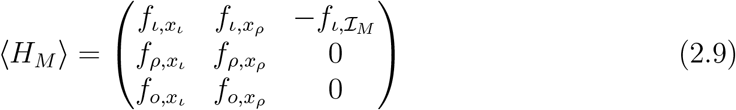

Here all partial derivatives 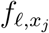 are evaluated at 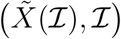.

#### Lemma 2.1.

*Let x*_*o*_(ℐ) *be input-output function of a multiple input parameter network. The partial derivative of x*_*o*_(ℐ) *with respect to the M-th parameter* ℐ_*M*_ *satisfies*

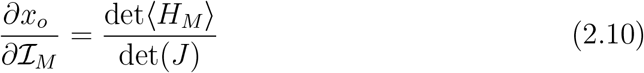

*Here* det(*J*) *and* det ⟨*H*_*M*_ ⟩ *are evaluated at the equilibrium point* 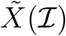. *Hence*,

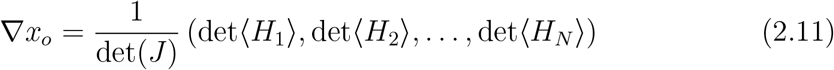

*Moreover*, ℐ^0^ *is a point of infinitesimal homeostasis if and only if*

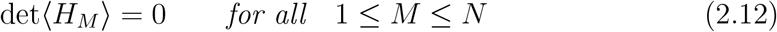

*as a function of* ℐ *evaluated at* ℐ^0^.

*Proof*. Implicit differentiation of the equation 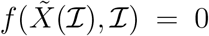 with respect to ℐ yields the linear system

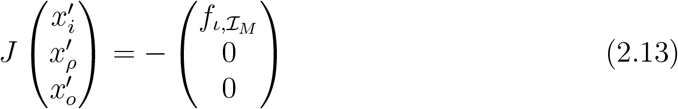

Since 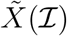 is assumed to be a linearly stable equilibrium, it follows that det(*J*) ≠ 0. On applying Cramer’s rule to (2.13) we can solve for 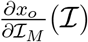 obtaining (2.10). Applying (2.10) to (2.7), we obtain equation (2.11). □

#### Remark 2.3.

An explicit expression for det ⟨*H*_*M*_⟩ can be obtained by expanding it with respect to the last column and the *ι*_*m*_-th row:

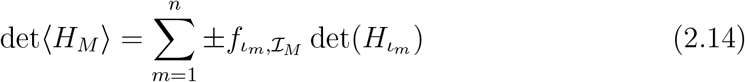

Here 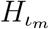 is obtained from *H* by removing the last column and the *ι*_*m*_-th row. When there is a single input parameter, i.e. *N* = 1, the gradient ∇*x*_*o*_ reduces to ordinary derivative 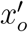 and (2.11) gives the formula for 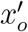 obtained in [38]. When there is a single input parameter and a single input node, *N* = *n* = 1, there is only one matrix 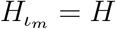, called the *homeostasis matrix* and (2.11) gives the corresponding formula for 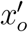 obtained in [55]. ◊

### 2.3 Core Networks

Wang et al. [55] introduced a fundamental construction for the study homeostasis in input-output networks, called ‘core subnetwork’. Madeira and Antoneli [38] extended this construction to the case of input-output networks with multiple input nodes and single input parameter. Here we extend the arguments of [38, 55] to the case of multiparameter networks.

Let (*X*\*, ℐ*) be a linearly stable equilibrium of the 𝒢-admissible ODE system (2.1). Then (*X*\*, ℐ*) it satisfies the system of equations

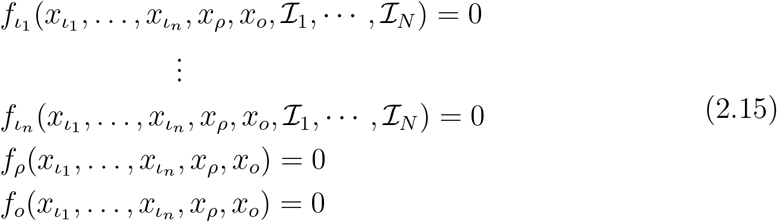

Partition the nodes of the network 𝒢 as follows: (i) input nodes (whose dynamics explicitly depends on at least one input), (ii) the output node and (iii) the regulatory nodes, that can be classified in three types depending if they are upstream from the output node or/and downstream from at least one input node. More precisely, the set of regulatory nodes may be partitioned as:

a. those nodes *σ* that are both upstream from *o* and downstream from at least one input node *ι*_*m*_,
b. those nodes *d* that are not downstream from any input node *ι*_*m*_,
c. those nodes *u* which are downstream from at least one input node *ι*_*m*_, but not upstream from *o*.

Figure 1 shows the types of connections which can be found in 𝒢.

**Figure 1:**
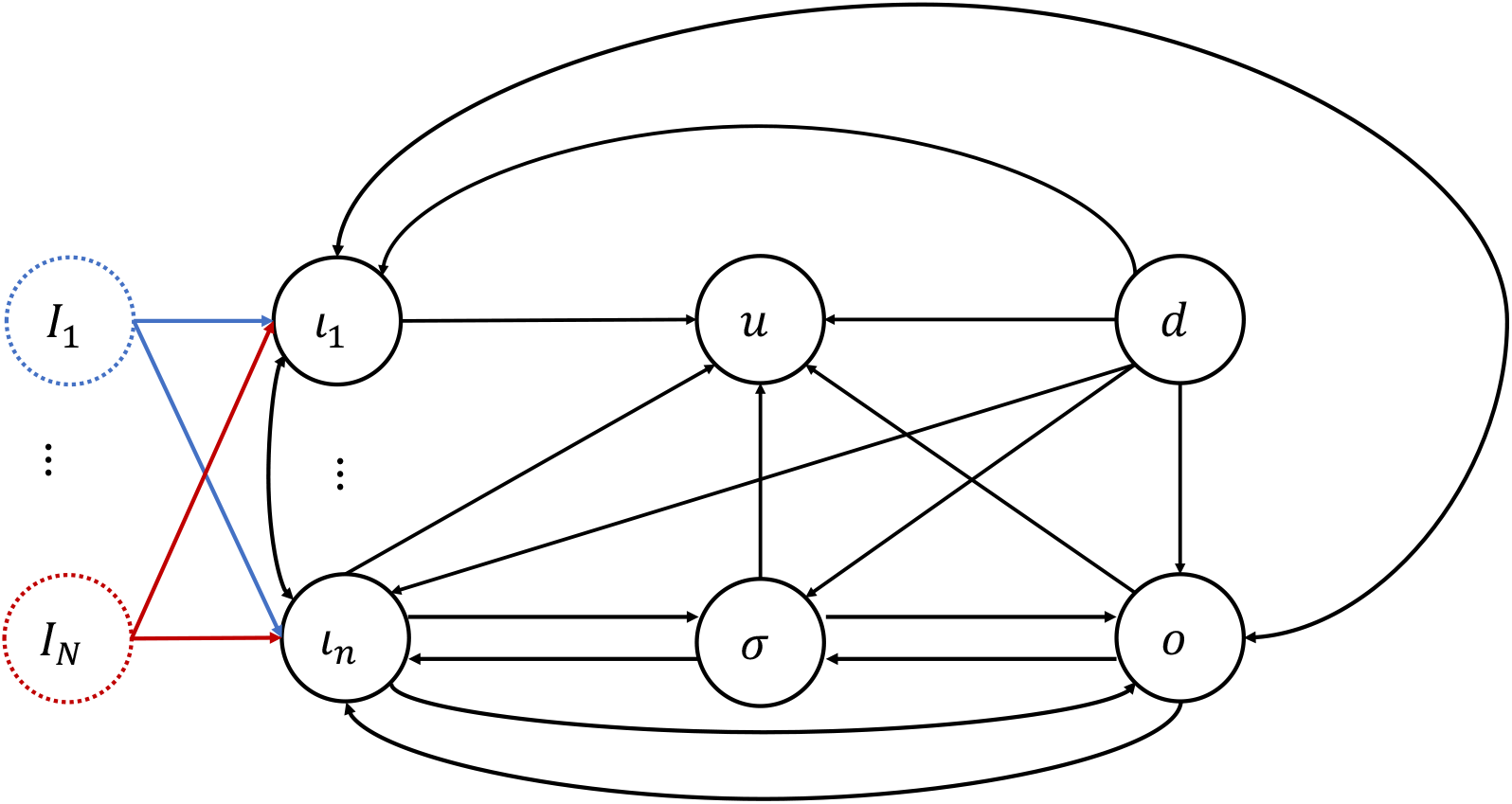
The possible connections in 𝒢.

#### Definition 2.4.

Let 𝒢 be a multiparameter input-output network. The *core subnetwork* 𝒢_*c*_ of 𝒢 is the subnetwork whose nodes are: (i) the input nodes *ι*_1_, …, *ι*_*n*_, (ii) the regulatory nodes *σ* that are upstream from the output node and downstream of at least one input node, and (iii) the output node *o*. The arrows of 𝒢_*c*_ are the arrows of 𝒢 connecting the nodes of 𝒢_*c*_.

#### Theorem 2.2.

*Let* 𝒢 *be a multiparameter input-output network and* 𝒢_*c*_ *the cor-responding core subnetwork. Then the input-output function* 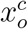 *of* 𝒢_*c*_ *exhibits in-finitesimal homeostasis at* ℐ* *if and only if the input-output function x*_*o*_ *of* ℐ *exhibits infinitesimal homeostasis at* ℐ*.

*Proof*. Follows from Theorem 3.2. □

Theorem 2.2 allows one to assume, without loss of generality, that 𝒢 is a *core network*, that is 𝒢 = 𝒢_*c*_, as far as infinitesimal homeostasis is concerned.

### 2.4 Structure of Infinitesimal Homoestasis

In this subsection, unless explicitly stated, we assume that 𝒢 is a core multiparameter input-output network with input nodes *ι*_1_, …, *ι*_*n*_ and input parameters ℐ_1_, …, ℐ_*N*_.

By Lemma 2.1 a network 𝒢 exhibits infinitesimal homeostasis at a point ℐ^0^ whenever the vector-valued function (when evaluated at 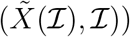 vanishes at ℐ^0^:

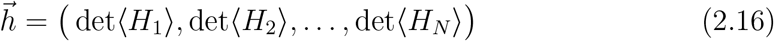

Here det ⟨*H*_*M*_ ⟩ are the determinants of the ℐ_*M*_ - generalized homeostasis matrices.

In order to analyze and simplify these determinants let us introduce some terminology. A *multivariate vector-valued polynomial*, or, simply a *polynomial mapping* is a mapping *P* : ℝ^*k*^ → ℝ^*k*^ with polynomial components. That is, if we write *P* in components as

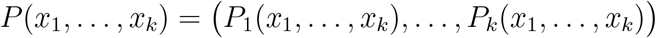

then *P*_*k*_ : ℝ^*k*^ → ℝ are multivariate polynomial functions. We say the *P* is irreducible if and only if each component *P*_*j*_ is irreducible. Suppose there is a multivariate polynomial function *p* : ℝ^*k*^ → ℝ that is common factor to all components *P*_*j*_. Then we can factor *p* from the polynomial vector *P* as

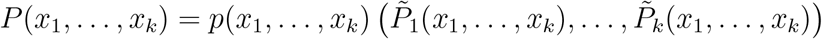

We say that *p* is a *scalar factor* of *P*.

Recall that the nonzero entries of the ℐ_*M*_ - generalized homeostasis matrices ⟨*H*_*M*_⟩ are the partial derivatives 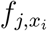 and 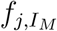. In particular, det ⟨*H*_*M*_⟩ is a homogeneous polynomial function of degree (*n* + *r* + 1) in the partial derivatives 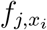 and 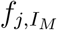. Hence, the vector-valued function 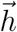 is a (formal) polynomial mapping on the ‘variables’ 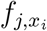 and 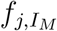. The scalar-valued function 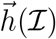 (depending on the multiparameter ℐ) is obtained by evaluating the partial derivatives 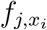 and 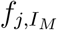 at 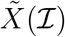.

Let us motivate the next definition with a simple observation. In the multiparamter case, even a core network have nodes that are not affected by some input parameters. For example, consider the 5-node multiparameter network 𝒢 shown in Figure 2. It has three input nodes, *ι*_1_, *ι*_2_ and *ι*_3_, and two input parameters, ℐ_1_ (blue) and ℐ_2_ (red). Although, 𝒢 is a core network, the input node *ι*_1_ is not affected by the parameter ℐ_2_, and the node *ι*_3_ is not affected by parameter ℐ_1_. To overcome this difficulty, we define the ‘specialized networks’ relative to a single input parameter.

**Figure 2:**
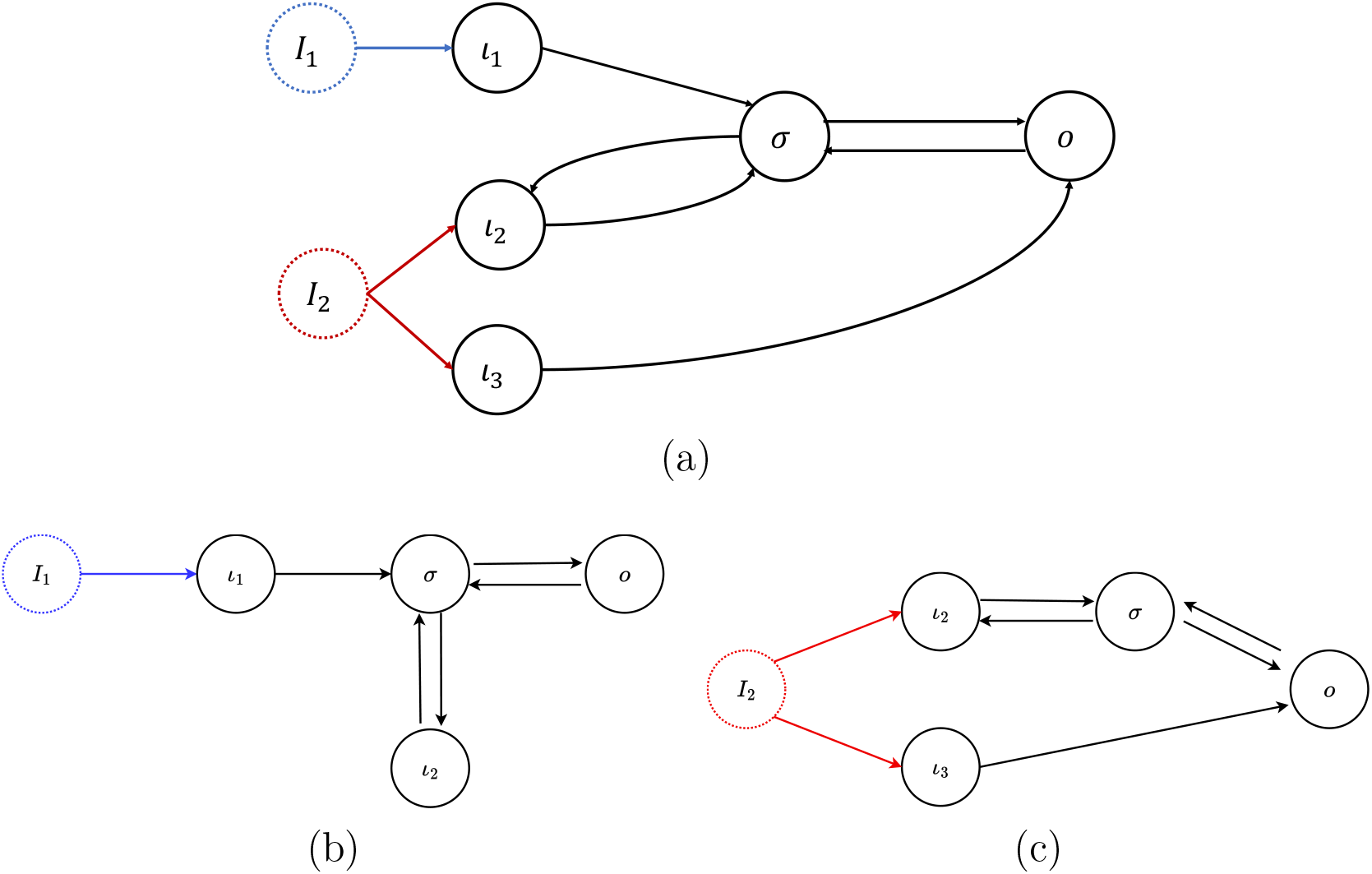
In (a) we show a 2-input parameter input-output network 𝒢, with 5 nodes (three input nodes *ι*_1_, *ι*_2_, *ι*_3_, one output node *o* and one regulatory node *σ*). In (b) and (c) we show the two *specialized subnetworks* of 𝒢, which are always single input parameter multiple input node networks (see Definition 2.5). The ℐ_1_-specialized subnetwork 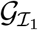, shown in (b), has 1 input node (*ι*_1_). Notice that *ι*_2_ is a regulatory node for this network. The ℐ_2_-specialized subnetwork 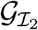, shown in (c), has 2 input nodes (*ι*_2_, *ι*_3_).

#### Definition 2.5.

Let 𝒢 be a core multiparameter network with input parameters ℐ_1_, …, ℐ_*N*_. The ℐ_*M*_ *-specialized subnetwork* 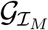 is the input-output subnetwork of 𝒢 consisting of all the input nodes that receive the input parameter ℐ_*M*_, all the regulatory nodes that are downstream from those input nodes and the output node. The arrows of 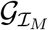 are the arrows of 𝒢 between the nodes of 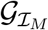. The subnetwork 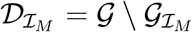 generated by the nodes in 𝒢 that do not belong to 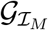 is called the ℐ_*M*_ *-vestigial subnetwork*.

The specialized subnetwork 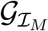 can be considered as a multiple input node network with single input parameter ℐ_*M*_, as studied in [38], by ‘forgetting’ the effect of the other parameters and reducing to the core network using the core reduction theorem of [38, Thm. 3.2]. The input nodes of 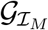 are exactly the input nodes of 𝒢 that are affected by the parameter ℐ_*M*_.

#### Definition 2.6.

Let 𝒢 be a core multiparameter network with input parameters ℐ_1_, …, ℐ_*N*_. The ℐ_*M*_ *-homeostasis matrix* 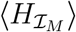 associated to 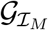, is the generalized homeostasis matrix of the multiple input-output network 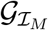 (see [38, Sec 2.3]). Similarly, the corresponding ℐ_*M*_ - vestigial subnetwork 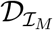 has an associated *jacobian matrix* 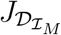(see [38, Sec 3.2]). To simplify the notation, in the case where the ℐ_*M*_ - vestigial subnetwork is empty 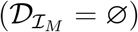, we write 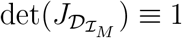.

Now we need to specify the set Ω ⊆ ℝ^*N*^ of allowable parameter values. This set depends on the admissible vector field *F* and the type of model being considered. For instance, in biochemical network models Ω is the positive orthant of ℝ^*N*^. The subset of *non-singular parameters* of *F* on Ω is defined as

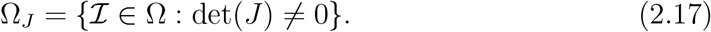

where 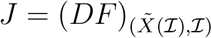 is the jacobian. The set Ω_*J*_ also depends on the vector field *F* and, generically, is an open dense subset of Ω.

#### Lemma 2.3.

*For each M* = 1, …, *N, we have:*

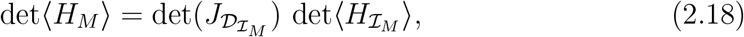

*Moreover*, 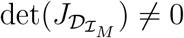 *over* Ω_*J*_ *and the irreducible factors of* 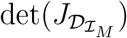 *never are irreducible scalar factors of* 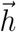.

*Proof*. In case 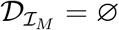, the result follows from the convention that 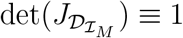. In case 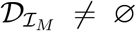, the vestigial subnetwork is composed by nodes that are not downstream from the input nodes affected by the parameter ℐ_*M*_. Hence, we can apply to 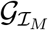 the ‘core network’ theorem for networks with multiple input nodes and a single input parameter [38, Thm 3.2]. The statement about the irreducible factors of 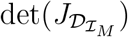 follows from an argument similar to the one in [38, Prop 3.8]. □

Lemma 2.3 allows us further simplify the components of 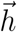, by considering the determinants 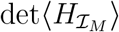. This reduce to the situation already studied in [38].

#### Definition 2.7.

The *vector determinant* associated to an input-output function *x*_*o*_ is the vector-valued function defined by

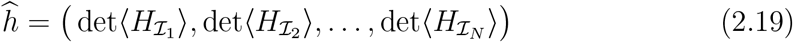

where 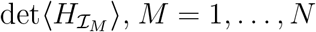, is the determinant of generalized homeostasis matrix of the ℐ_*M*_ - specialized subnetwork 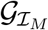. In particular, we can consider the vector-valued function 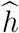 as a (formal) polynomial mapping on the ‘variables’ 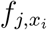 and 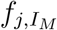.

#### Proposition 2.4.

*The vector-valued functions* ∇*x*_*o*_, 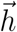 *and* 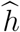, *defined on* Ω → ℝ^*N*^, *have the same set of zeros on* Ω_*J*_.

*Proof*. The first equality follows from Lemma 2.1 and the second equality follows from Lemma 2.3. □

The König-Frobenius theorem [10,49] (see also [38,55]) imply that the components of the polynomial mapping 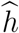 can be factorized as the product of the determinants of the irreducible diagonal blocks of each 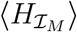 (defined up to row and column permutations).

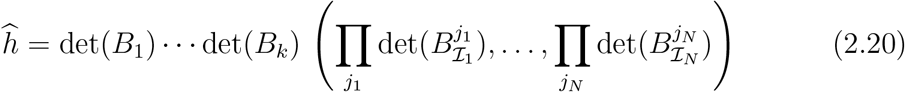

where the scalar factors det(*B*_*j*_) (*j* = 1, …, *k*) are such that the matrices *B*_*j*_ are common irreducible diagonal blocks of all matrices 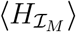. Therefore, we can split the problem of classifying homeostasis types supported by 𝒢 into two cases according to whether the components of 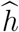 is a common scalar factor or not.

#### Definition 2.8.

Let 𝒢 be a core multiparameter network. An irreducible block *B* of some 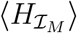 is called a *homeostasis block* A homeostasis block *B* such that det(*B*) is a common scalar factor of 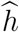 is called *pleiotropic*. The other homeostasis blocks of are called *coincidental*.

Recall that the *homeostasis types* of a single parameter input-output network 𝒢 is given in terms of the factors of *h* = det(*H*) (see [38, 55])

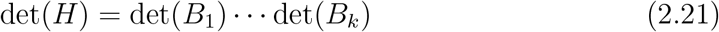

where each irreducible block *B*_*j*_ can be of three types, called *appendage, structural* and *counterweight*. There is only one counterweight block defined as the only irreducible block that contains all the partial derivatives of *f* with respect to the input parameter. Infinitesimal homeostasis of type *B*_*j*_ occurs if det(*B*_*j*_) = 0 and det(*B*_*i*_) ≠ 0 for all *i* ≠ *j*. This is generic when there is only one input parameter.

Now, suppose that 𝒢 has *N* input parameters affecting *n* input nodes. Generically, it is expected that *N* irreducible factors of 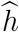 can simultaneously vanish at a fixed input parameter value.

#### Definition 2.9.

Let 𝒢 be a multiparameter core network. Let *B* be a homeostasis block of size *ℓ*. We say that the *homeostasis class* of *B* is

a. *input counterweight* if *B* contains partial derivatives with respect the input parameter,
b. *appendage* if *B* has *ℓ* self-couplings,
c. *structural* if *B* has exactly *ℓ* − 1 self-couplings.

It follows from the argument in [38, Sec.3.4] applied to the specialized subnetowrks 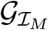, that each homeostasis block of 𝒢 is of one of the classes in Definition 2.9.

#### Definition 2.10.

Let 𝒢 be a core multiparameter network.

a. We say that *pleiotropic homeostasis* occurs when at least one pleiotropic block has vanishing determinant at some fixed input parameter value. The pleiotropic blocks determine the *pleiotropic homeostasis types* of 𝒢.
b. We say that *coincidental homeostasis* occurs when a *N* - upla of coincidental blocks 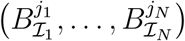 has simultaneously vanishing determinants at some fixed input parameter value. The *N* - uplas of coincidental blocks determine the *coincidental homeostasis types* of 𝒢.

#### Definition 2.11.

Let 𝒢 be a core multiparameter network and *B* be a homeostasis block. The *homeostasis subnetwork* 𝒦_*B*_ associated to *B* is defined as follows. The nodes of 𝒦_*B*_ are the nodes *σ* and *ρ* of 𝒢 such that 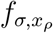 is a non-zero entry of *B*. The arrows of 𝒢_*B*_ are the arrows *σ* → *ρ* of 𝒢 such that *σ, ρ* ∈ 𝒦_*B*_ with *σ* ≠ *ρ*.

### 2.5 Pleiotropic Homeostasis Types

In this section we classify the pleiotropic sub-blocks of a multiparameter core network.

#### Proposition 2.5.

*Let* 𝒢 *be a multiparameter core network and B be a pleiotropic block of* 𝒢.

i. *Then B is either appendage or structural*.
ii. *More precisely, B is an appendage (respect. structural) block if and only if it is* 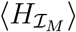*-appendage (respect*. 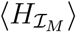*-structural) block, for all M* = 1, …, *N*.

*Proof*. (i) From the results of [38, 55], we see that, for each *M* = 1, …, *N, B* can be classified with respect to the specialized subnetwork 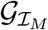 associated to the input ℐ_*M*_ as an 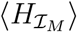-input counterweight, an 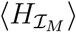-structural or an 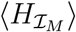-appendage block. As the derivatives 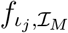 would appear in the expression of det(*B*) if it was a 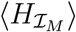-input counterweight block, we conclude that det(*B*) must be either an 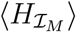-structural or an 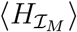-appendage block, for all *M* = 1, …, *N*.

(ii) This follows immediately from the fact that the classification is based on the number of self-coupling entries. □

In remark 2.3 we observed that when the network 𝒢 has a single parameter the theory developed here reduces to the situation considered in [38]. The next result shows that when the multiparameter network 𝒢 has a single input node then the theory essentially reduces to the case where there is only one input parameter, considered in [55]. This is an extreme case where only pleiotropic homeostasis occurs.

#### Proposition 2.6.

*Suppose the core multiparameter network* 𝒢 *has only one input node ι and multiple input parameters* ℐ_1_, …, ℐ_*N*_. *Then, we have:*

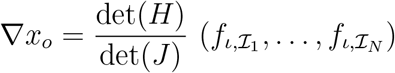

*where* det(*H*) *is the homeostasis determinant of the network* 𝒢, *as a polynomial function of the partial derivatives* 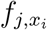. *In particular, condition* (2.6) *implies that infinitesimal homeostasis occurs if and only if* det(*H*) = 0.

*Proof*. This is a consequence of Lemma 2.1 and of Equation 2.14 for det⟨*H*_*M*_ ⟩ when 𝒢 has a single input node. □

Therefore, Proposition 2.6 implies that the classification of homeostasis types of a multiparameter network with a single input node is exactly the same as that of a single input parameter single input node network. However, it is not true that we expect to see the same ‘homeostasis phenomena’ in both networks.

Now, suppose that 𝒢 has a single input node, but multiple input parameters (ℐ_1_, …, ℐ_*N*_) affecting the input node. Consequently, the function *h*, which is the same polynomial on the partial derivatives 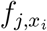, becomes a multivariate function when evaluated at 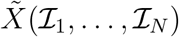. Generically, it is expected that *N* irreducible factors of (2.21) can simultaneously vanish at a fixed 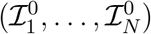. This is very similar to the kind of ‘homeostasis mode interaction’ considered in [13], although in this paper the multiparameter does not necessarily represent an external stimulus, but can represent a node self-coupling strength or a coupling strength.

Returning to the general case, the next result gives a complete topological classification of the homeostasis subnetworks corresponding to the pleiotropic homeostasis types.

#### Theorem 2.7.

*Let* 𝒢 *be a multiparameter core network, B be a pleiotropic block of* 𝒢 *and* 𝒦_*B*_ *be the corresponding homeostasis subnetwork to B in* 𝒢.

i. *Then* 𝒦_*B*_ *is either an appendage or structural subnetwork of all* ℐ_*M*_ *-specialized subnetworks* 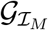.
ii. *More precisely*, 𝒦_*B*_ *is an appendage (respect. structural) subnetwork if and only if it is a* 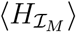*-appendage (respect*. 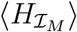*-structural) subnetwork, for some (and hence for all) M* = 1, …, *N*.

*Proof*. The result follows from Theorems 3.3 and 3.4 for the appendage case and Theorems 3.8 and 3.9 for the structural case. See section 3.2 for precise characterization of each type of subnetwork. □

### 2.6 Coincidental Homeostasis Types

The occurrence of coincidental homeostasis reflects the fact that the mechanism leading to homeostasis is not the same with respect all the input parameters. The coincidental homeostasis types are given by all the possible combinations of coincidental blocks. A coincidental type can be of structural class, appendage class or input counterweight class. Thus, depending on the network, a coincidental type can have the form of a mix of these three classes.

In the simplest case the determinant of a coincidental block can be an entry of some 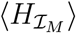 of the form 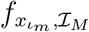. Since, by assumption, these entries cannot vanish it may happen that some coincidental types do not yield infinitesimal homeostasis.

When we have a *proper coincidental type* (i.e., all determinants can vanish) there is the possibility of non-trivial interaction between the homeostasis subnetworks with respect to each other. This feature gives an astonishing array of phenomena that can arise even in small networks.

In Figure 3 we show three examples of small 2-parameter core networks with very distinct behavior. These examples are discussed below.

**Figure 3:**
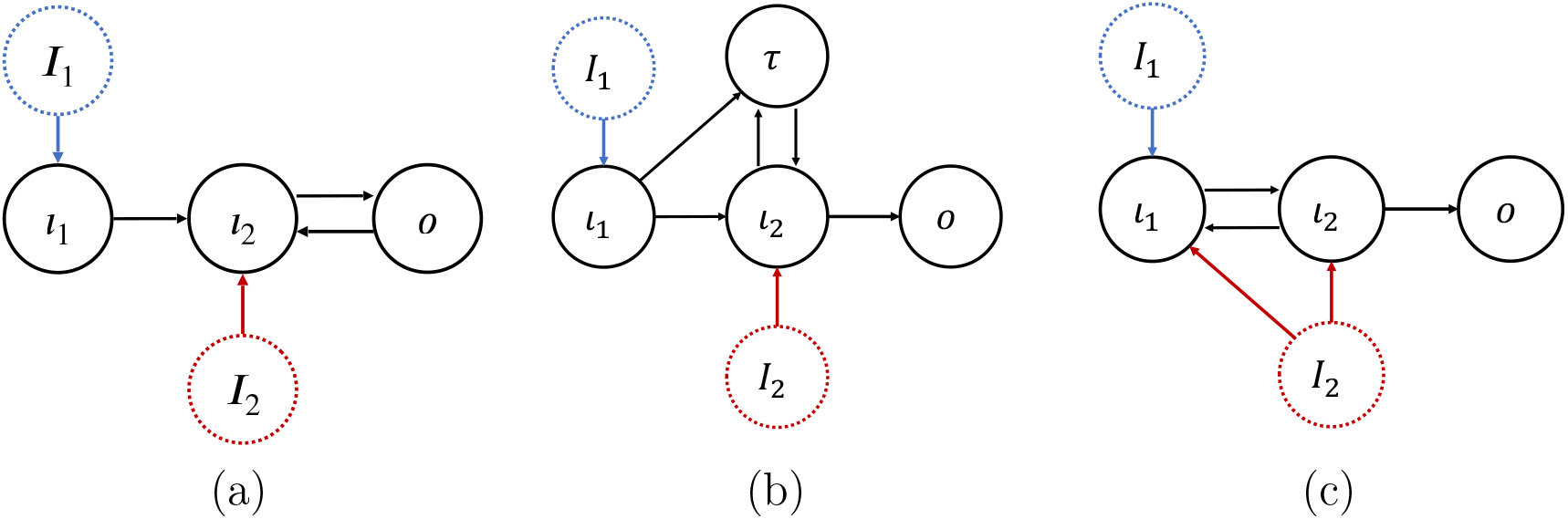
Core networks with two input nodes *ι*_1_ and *ι*_2_ and two input parameters ℐ_1_ and ℐ_2_. All three networks have pleiotropic-structural subnetwork of Haldane type given by *ι*_2_ → *o*. Regarding coincidental homeostasis: (a) Has 2 coincidental types but none of them is proper (see Example 2.12). (b) Has 4 coincidental types but only one of the form (structural, appendage) is proper (see Example 2.13). (c) Has 4 coincidental types but only one of the form (structural, coincidental) is proper. However the simultaneous vanishing of these determinants implies the vanishing of the jacobian determinant, as well. That is, the simultaneous occurrence of critical point and bifurcation (see Example 2.14).

#### Example 2.12.

Consider the 2-parameter core network 𝒢 shown in Figure 3(a). For such network, one has

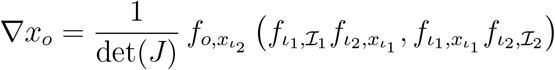

where 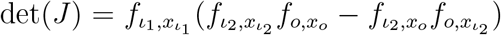. The specialized subnetwork 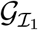 and the specialized subnetwork 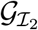 are, respectively,

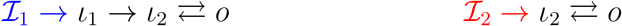

Therefore, the vector determinant is given by

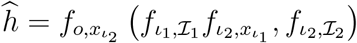

Note that, by definition, 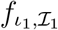 and 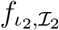 cannot vanish. Hence, the network does not support coincidental homeostasis, only pleiotropic-structural, given by the vanishing of 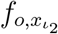. Here, the obstruction to coincidental homeostasis is due to the fact that the second component of *ĥ* is a non-vanishing counterweight factor 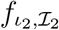. ◊

#### Example 2.13.

Consider the 2-parameter core network 𝒢 shown in Figure 3(b). For such network one has

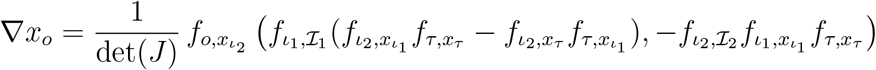

where 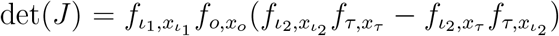. The specialized subnetwork 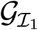 is the same as the full network 𝒢 without the input parameter ℐ_2_ at node *ι*_2_ and the specialized subnetwork 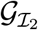 is given by

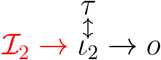

Therefore, the vector determinant is given by

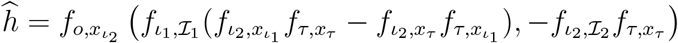

Pleiotropic homeostasis can occur by the vanishing of 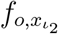. Here, the input node *ι*_2_ associated to the parameter ℐ_2_ is a super-simple node for both specialized subnetworks. But, unlike Example 2.12, coincidental homeostasis can occur here. In order for coincidental homeostasis to occur both irreducible determinants below must vanish simultaneously

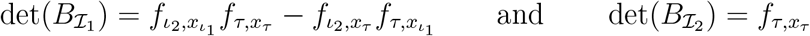

Here, 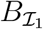 is the coincidental block associated to the structural (feedforward) subnetwork generated by {*ι*_1_, *ι*_2_, *τ*} and 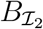 is the coincidental block associated to the appendage (null-degradation) subnetwork generated by {*τ*}. When 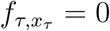 vanishes it reduces the vanishing of 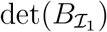 to the vanishing of either 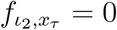 or 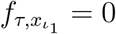, generically. On one hand, if 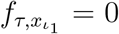 then det(*J*) ≠ 0 and we have the occurrence coincidental homeostasis. On the other hand, if 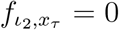 then det(*J*) = 0 simultaneously. Thus a steady-state bifurcation occurs exactly at the critical (homeostasis) point. ◊

#### Example 2.14.

Consider the 2-parameter core network 𝒢 shown in Figure 3(c). For such network one has

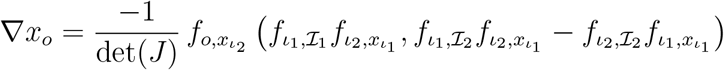

where 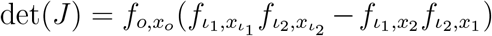. The specialized subnetwork 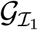 is the same as the full network 𝒢 without the input parameter ℐ_2_ at nodes *ι*_1_ and *ι*_2_ and the specialized subnetwork 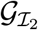 is the same as the full network 𝒢 without the input parameter ℐ_1_ at node *ι*_1_. Therefore, the vector determinant is given by

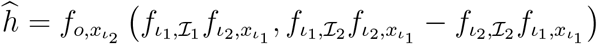

Pleiotropic homeostasis can occur by the vanishing of 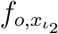. In order for coincidental homeostasis to occur both irreducible determinants below must vanish simultaneously

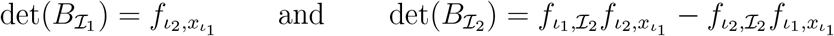

Here, 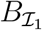 is the coincidental block associated to the structural (Haldane) subnetwork generated by {*ι*_1_, *ι*_2_} and 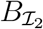 is the coincidental block associated to the counterweight subnetwork generated by {*ι*_1_, *ι*_2_}. Hence, occurrence of coincidental homeostasis requires that 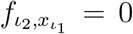 and 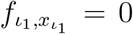. But this implies that det(*J*) = 0. That is, steady-state bifurcation occurs exactly at the critical (homeostasis) point. Unlike Example 2.13, there is no instance where only homeostasis occurs. ◊

In Examples 2.13 and 2.14 we found the simultaneous occurrence of steady-state bifurcation with the critical point. This is a kind of “mode interaction” between coincidental blocks that is distinct from the one discussed in [13], where the mode interaction is caused by the vanishing of multiple pleiotropic blocks. It seems that this interaction is ‘caused’ by the overlapping between the subnetworks associated to coincidental blocks in distinct components of the vector determinant.

The of infinitesimal homeostasis (Definition 2.1) excludes the simultaneous occurrence of steady-state bifurcation with the critical point. Hence, strictly speaking, Example 2.14 does not supports coincidental homeostasis. However, if one considers extending the definition of homeostasis to include such cases (see [14, 15] for some advances in this direction) then one may get a much richer variety of phenomena. For instance, in Example 2.13 coincidental homeostasis associated to the same pair of coincidental blocks has two “flavors” according to which factor vanishes.

As we have seen there are networks that do support both pleiotropic and coincidental homeostasis (e.g., the network of Example 2.13) and some networks that support only one of each type. For instance, the network of Example 2.12 supports only pleiotropic homeostasis. More generally, as shown in Proposition 2.6, networks that have only one input node, only support pleiotropic homeostasis. It is easy to find examples of networks that do not support only coincidental homeostasis.

Still, one may wonder whether there is a multiparameter core network that do not support either of them, that is, it does not support infinitesimal homeostasis. The next proposition shows that this cannot happen, namely, any multiparameter core network always support at least one type of homeostasis.

#### Proposition 2.8.

*A multiparameter core network* 𝒢 *always supports infinitesimal homeostasis*.

*Proof*. In order to prove the proposition, consider all input nodes *ι*_*m*_, such there is an *ι*_*m*_-simple node *σ* (*σ* ≠ *ι*_*m*_) that receives an arrow from *ι*_*m*_ and such that 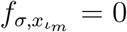 (Haldane homeostasis). Let 𝒢_*m*_ be the core subnetwork between the input node *ι*_*m*_ and output node *o* [38, Def. 2.13]. By definition, 𝒢_*m*_ is a single input-output network. Then, by the characterization of structural homeostasis in networks with single one input node (see [55]), the homeostasis determinant of 𝒢_*m*_ vanish, i.e., if 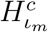 is the homeostasis matrix of 𝒢_*m*_, then det 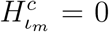. This fact together with [38, Eq 3.39] implies that 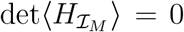, for all *M* = 1, …, *N*. To conclude the argument, we claim that this construction does not force det(*J*) = 0, generically. This follows from that fact that one of the terms that appear in the expression of the Jacobian determinant is the product of all the self-couplings of nodes of 𝒢. As the construction above does not assume that the self-couplings to be equal to zero, the claim holds. □

#### Remark 2.15.

It seems tempting to say that the ‘pleoitropic part’ of a multiparameter core network 𝒢 is given by a single input subnetwork that generated by the nodes that are ‘downstream’ from the first absolutely super simple node of 𝒢. Hence, by complementarity the ‘coincidenal part’ is given by the subnetwork generated by the nodes that are ‘upstream’ from the first absolutely super simple node of 𝒢. The problem with this putative characterization is the presence of *backward arrows*, which can be defined a the arrows of 𝒢 that correspond to terms 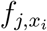 that do not belong to any irreducible block. Note that this definition of backward arrows given above is more general than the one given in [55], nevertheless, it implies that the removal of backward arrows does not alter the vector determinant *ĥ*. It may happen that some of these backward arrows connect nodes in the ‘pleoitropic part’ to nodes in the ‘coincidenal part’. We conjecture that once all the backward arrows are removed then there is a decomposition into ‘pleoitropic part’ and ‘coincidenal part’. ◊

### 2.7 Algorithm to Determine All Homeostasis Types

Using the results obtained here, together with [38, 55], one can write down a general algorithm to find all homeostasis types of a given multiparameter core network 𝒢.

**Step 1:** For each parameter ℐ_*M*_ with *M* = 1, …, *N* determine the ℐ_*M*_ - specialized subnetwork 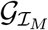 as in Definition 2.5.

**Step 2:** Since each 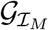 is a single parameter multiple input-output network one can apply the algorithm of [38, Sec. 2.6] to determine all the homeostasis subnetworks 𝒦_*B*_ of each 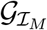. The corresponding irreducible factors det(*B*) can be computed directly from the homeostasis subnetworks 𝒦_*B*_.

**Step 3:** Determine the homeostasis subnetworks that are common to all 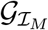. These homeostasis subnetworks correspond to the pleiotropic homeostasis types.

**Step 4:** The *N* - uplas formed by combinations of the remaining homeostasis subnetworks correspond to the coincidental homeostasis types.

## 3 Classification of Homeostasis Types

In this section we present the proofs of the main results of the paper.

### 3.1 Reduction to the Core Network

In this section, unless explicitly stated, we assume that 𝒢 is a multiparameter input-output network with input nodes *ι*_1_, …, *ι*_*n*_, and input parameters ℐ_1_, …, ℐ_*N*_.

The definition of core subnetwork implies that the admissible system of equations 2.15 can be written as

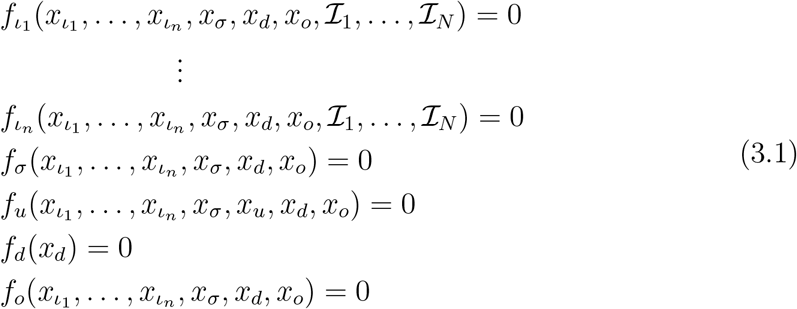

Now we can freeze the variables *x*_*d*_ at an appropriate value and obtain an admissible system for 𝒢_*c*_ from system (3.1).

#### Lemma 3.1.

*Suppose that the point* 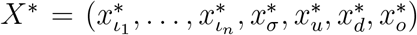 *is a linearly stable equilibrium of* (3.1). *Then the* 𝒢_*c*_*-admissible system obtained from* (3.1) *by freezing x*_*d*_ *at* 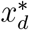, *given by*

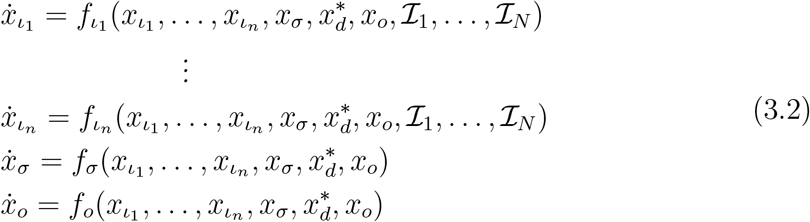

*has a linearly stable equilibrium* 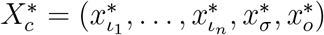.

*Proof*. It is trivial that 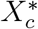 is an equilibrium of (3.2). As shown in [38, Lem 3.1], 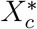 is linearly stable. Indeed, the Jacobian matrix *J* of (3.1) evaluated at *X*\* is

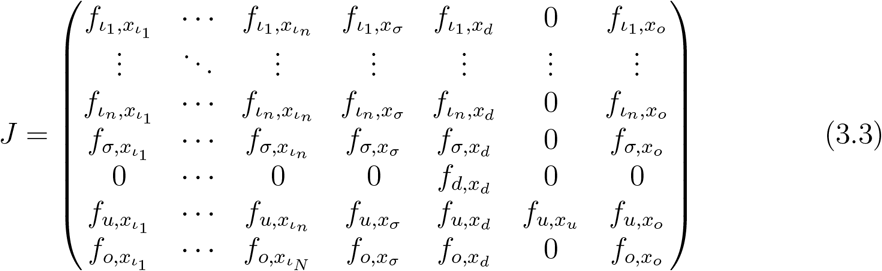

and therefore the eigenvalues of *J* are the same eigenvalues of 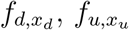 and of the matrix *J*_*c*_, where

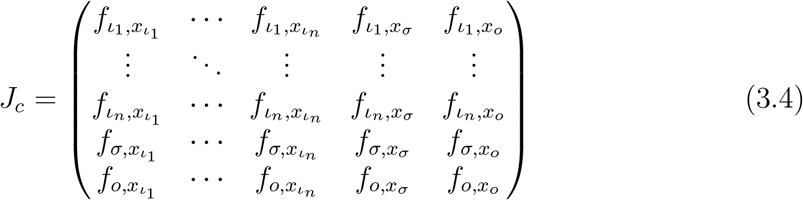

Since *J*_*c*_ is the Jacobian matrix of (3.2) calculated at 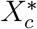, it follows that if *X*\* is a linearly stable equilibrium then so it is 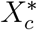. □

##### Theorem 3.2.

*Let x*_*o*_(ℐ) *be the input-output function of the admissible system 2*.*15 for the network* 𝒢 *and let* 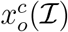 *be the input-output function of the admissible system* (3.2) *for the corresponding core network* 𝒢 _*c*_. *Then* 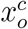 *exhibits infinitesimal homeostasis at* ℐ* *if and only if x*_*o*_ *exhibits infinitesimal homeostasis at* ℐ*.

*Proof*. For each weighted homeostasis matrix ⟨*H*_*M*_ ⟩, we have:

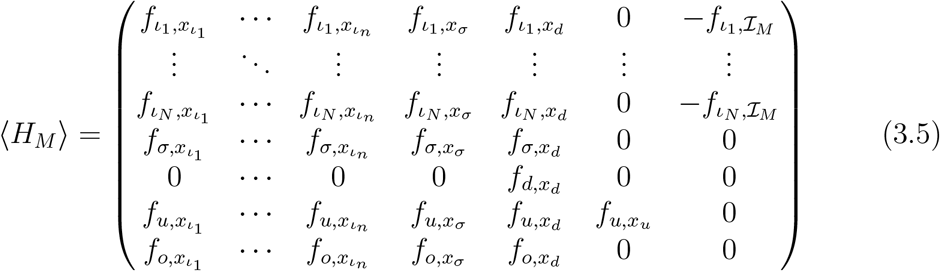

Hence, for each 1 ≤ *M* ≤ *N*, we have:

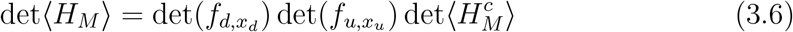

where

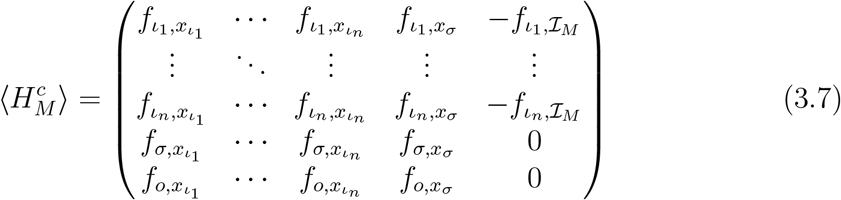

From Lemma 3.1, we have

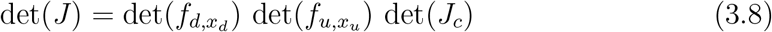

Applying (3.6) and (3.8) to (2.11), we get:

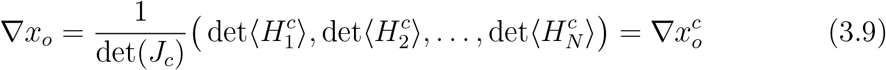

Therefore, *x*_*o*_ and 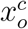 have exactly the same critical points. □

### 3.2 Classification of Pleiotropic Homeostasis Types

In this sub-section, unless explicitly stated, we assume that 𝒢 is a core multiparameter input-output network with input nodes *ι*_1_, …, *ι*_*n*_, and input parameters ℐ_1_, …, ℐ_*N*_, with *n, N* ≥ 2.

We shall now study the subnetworks associated to pleiotropic homeostasis blocks. Bearing this in mind, we start by extending the classification of nodes from [38].

#### Definition 3.1.

Let 𝒢 be a multiparamete core network.

a. A directed path connecting nodes *ρ* and *τ* is called a *simple path* if it visits each node on the path at most once.
b. An *ι*_*m*_*o-simple path* is a simple path connecting the input node *ι*_*m*_ to the output node *o*.
c. A node is *ι*_*m*_*-simple* if it lies on an *ι*_*m*_*o*-simple path.
d. A node is *ι*_*m*_*-appendage* if it is downstream from *ι*_*m*_ and it is not an *ι*_*m*_-simple node.
e. A node is ℐ_*M*_ - *absolutely simple* if it is an *ι*_*m*_-simple node, for every *m* such that 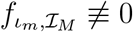.
f. A node is ℐ_*M*_ - *absolutely appendage* if it is an *ι*_*m*_-appendage node, for every *m* such that 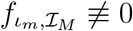.
g. An *ι*_*m*_-*super-simple node* is an *ι*_*m*_-simple node that lies on every *ι*_*m*_*o*-simple path.
h. An ℐ_*M*_ - *absolutely super-simple node* is a node that lies on every *ι*_*m*_*o*-simple path, for every *m* such that 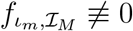.

It is immediate that the output node *o* is an ℐ_*M*_ - absolutely super-simple node, for all *M* = 1, …, *N*.

#### 3.2.1 Pleiotropic-Appendage Homeostasis

To study the structure of pleiotropic-appendage homeostasis, we shall first generalize the concepts of path equivalence and appendage subnetworks employed in [38, 55] to the current context.

##### Definition 3.2.

Let 𝒦 be a nonempty subnetwork of 𝒢. We say that nodes *ρ*_*i*_, *ρ*_*j*_ of 𝒦 are *path equivalent in* 𝒦 (or 𝒦-*path equivalent*) if there are paths in 𝒦 from *ρ*_*i*_ to *ρ*_*j*_ and from *ρ*_*j*_ to *ρ*_*i*_. A 𝒦*-path component* is a path equivalence class in 𝒦.

##### Definition 3.3.

The 𝒢*-complementary subnetwork* of an *ι*_*m*_*o*-simple path *S* is the subnetwork *CS* consisting of all nodes of 𝒢 not on *S* and all arrows in 𝒢 connecting those nodes.

##### Definition 3.4.

Let 𝒢 be a multiparamete core network.

a. For every *m* = 1, …, *n*, we define the *ι*_*m*_-*appendage subnetwork* 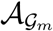 as the subnetwork of 𝒢 composed by all *ι*_*m*_-appendage nodes and all arrows 𝒢in connecting *ι*_*m*_-appendage nodes.
b. For every *M* = 1, …, *N*, we define the ℐ_*M*_ - *appendage subnetwork* 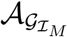 as the subnetwork of 𝒢 composed by all ℐ_*M*_ - absolutely appendage nodes and all arrows in 𝒢 connecting ℐ_*M*_ - absolutely appendage nodes. Note that

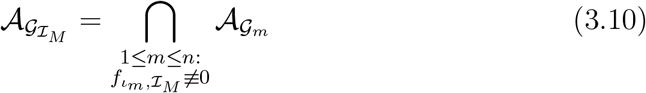
c. The *appendage subnetwork* 𝒜_𝒢_ is the subnetwork of 𝒢 composed by nodes which are ℐ_*M*_ - absolutely appendage, for all *M* = 1, …, *N*, and the arrows connecting such nodes. Note that

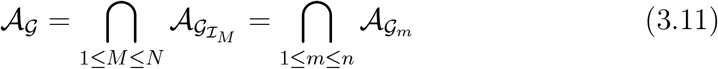

Now we can characterise the structure of pleiotropic-appendage homeostasis. Let *B* be a pleiotropic appendage block. By a similar argument employed in [38], we conclude that *B* must be the jacobian matrix of the corresponding subnetwork 𝒦_*B*_.

##### Theorem 3.3.

*Let* 𝒦 _*B*_ *be a subnetwork of* 𝒢 *associated with a pleiotropic-appendage block B. Then the following statements are valid:*

i. *Each node in* 𝒦_*B*_ *is an I*_*M*_ *-absolutely appendage node, for all M* = 1, …, *N*.
ii. *For every ι*_*m*_*o-simple path S, nodes in* 𝒦_*B*_ *are not CS-path equivalent to any node in CS* \ 𝒦_*B*_, *for all m* = 1, …, *n;*
iii. 𝒦_*B*_ *is a path component of* 𝒜 _𝒢_.

*Proof*. Statements (*a*) and (*b*) follow by applying [38, Thm 3.11] to each of the core subnetworks 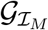. Statement (*c*) is proved along the same line as [38, Thm 3.11c]. □

Now we shall verify that the conditions listed in theorem 3.3 are also sufficient to guarantee the existence of a pleiotropic-appendage homeostasis block.

##### Theorem 3.4.

*Suppose* 𝒦 *is a subnetwork of* 𝒢 *such that:*

i. 𝒦 *is an* 𝒜 _𝒢_*-path component;*
ii. *For every ι*_*m*_*o-simple path S, nodes in* 𝒦 *are not CS-path equivalent to any node in CS* \ 𝒦_*j*_, *for all m* = 1, …, *n*.

*Then* det(*J*_𝒦_) *is an irreducible factor of* 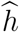.

*Proof*. Apply [38, Thm 3.13] to each of the specialized subnetworks 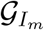. The validity of condition (*b*) of [38, Thm 3.13] for each specialized subnetwork 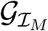 follows directly of condition (*b*) of this theorem. It is then enough to prove that 𝒦 is a pnath component of 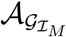, for all *m* = 1, …, *n*. As 𝒦_*j*_ is a path component of 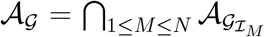 then for each *M* = 1, …, *N*, there is a 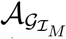 - path component 𝒯_*M*_ such that 𝒦 ⊆ 𝒯_*M*_. By condition (*b*), it follows that 𝒦 = 𝒯_*M*_, for each *M* = 1, …, *N*. □

#### 3.2.2 Pleiotropic-Structural Homeostasis

Now we shall study the pleiotropic-structural blocks.

Let 𝒱^𝒢^ be the set of nodes of 𝒢, 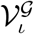 the set of nodes that are *ι*_*m*_-super simple, for all *m* = 1, …, *n* and 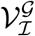 the set of nodes that are ℐ_*m*_-absolutely super-simple, for all *M* = 1, …, *N*. In [38], we introduced the notion of absolutely super-simple nodes with respect to the input nodes. This suggests that we can define absolutely super-simple nodes with respect to the input parameters. This leads to the question: Which subset of 𝒱^𝒢^ is more suitable to base the characterization of pleiotropic-structural subnetworks: 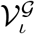 or 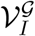. The simple, yet paramount, observation that the answer to this question is that both sets are equal.

##### Lemma 3.5.

*Let* 𝒢 *be a multiparamete core network. Then* 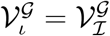.

*Proof*. It is enough to verify that 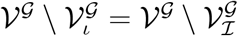. First, suppose there is a node 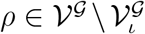. Then, there is at least one input node *ι*_*m*_ such that *ρ* is not an *ι*_*m*_-super-simple node. As 𝒢 is a core network, there is an input ℐ_*M*_ such that 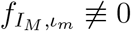, which implies that *ρ* is not ℐ_*M*_-absolutely super-simple and hence 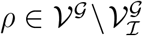. On the other hand, if 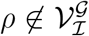, then there exists *M* such that *ρ* is not ℐ_*M*_ - absolutely super-simple ⇒ *ρ* is not *ι*_*m*_-super-simple, for some *m* such that 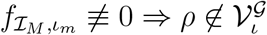. □

The importance of Lemma 3.5 is that it allows us to study the set 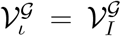 through either the characterization with respect to the input nodes or to the input parameters, whichever is more convenient. In particular, we can easily extend many of the results obtained in [38].

A slightly modification of the argument of Lemma 3.5 shows that the set of nodes that are *ι*_*m*_-simple, for all *m* = 1, …, *n*, and the set of nodes ℐ_*M*_ - absolutely simple, for all *M* = 1, …, *N*, are also equal. These observations justify the generalization of the concept of absolutely simple and absolutely super-simple nodes.

##### Definition 3.5.

Let 𝒢 be a multiparamete core network.

a. A node *ρ* is called *absolutely super-simple* if and only if it is an *ι*_*m*_-super simple node, for all *m* = 1, …, *n*. Equivalently, *ρ* is called *absolutely super-simple* if and only if it is an *I*_*M*_ - absolutely super-simple node, for all *M* = 1, …, *N*.
b. A node *ρ* is called *absolutely simple* if and only if it is an *ι*_*m*_ −simple node, for all *m* = 1, …, *n*. Equivalently, *ρ* is called *absolutely simple* if and only if it is an *I*_*M*_ - absolutely simple node, for all *M* = 1, …, *N*.

Following [38], we now define a total order relation in the set of absolutely super-simple nodes.

##### Definition 3.6.

Let 𝒢 be a multiparamete core network. Define a relation on 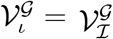 as follows: for any pair of nodes *σ*, 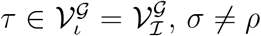 we write *σ* > *ρ* when *ρ* is downstream from *σ* by all *ι*_*m*_*o*-simple paths, for any *m* = 1, …, *n*.

##### Lemma 3.6.

*The relation on* 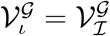 *given in Definition 3.6 is a total order*. □

*Proof*. This result is analogous to [38, Lem 3.15]

Consider now the ordered elements of 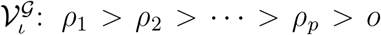. Similarly to [38, 55], we say that two elements *ρ*_*k*_ *> ρ*_*k*+1_ of 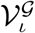 are *adjacent* when *ρ*_*k*+1_ is the first element of 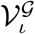 which appears after *ρ*_*k*_ in that ordering. We can now use this concept to introduce the elements that are crucial to characterise pleiotropic-structural homeostasis blocks.

##### Definition 3.7.

Let *ρ*_*k*_ *> ρ*_*k*+1_ be adjacent elements of 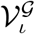. An *ι*_*m*_-absolutely simple node *ρ* is *between ρ*_*k*_ and *ρ*_*k*+1_ if there exists an *ι*_*w*_*o*-simple path that includes *ρ*_*k*_ to *ρ* to *ρ*_*k*+1_ in that order, for some *w* = 1, …, *n*.

The idea is to construct the structural subnetworks employing the concepts above, as it was done in [38, 55].

The *absolutely super-simple subnetwork*, denoted ℒ(*ρ*_*k*_, *ρ*_*k*+1_), is the subnetwork whose nodes are absolutely simple nodes between *ρ*_*k*_ and *ρ*_*k*+1_ and whose arrows are arrows of 𝒢 connecting nodes in ℒ (*ρ*_*k*_, *ρ*_*k*+1_). As we can characterise the absolutely super-simple and absolutely simple nodes (and consequently the absolutely super-simple subnetworks) with respect to each input node, we can construct the basic unit of pleiotropic-structural homeostasis in the same way the basic unit of structural homeostasis was constructed in [38].

##### Definition 3.8.

Let *ρ*_*k*_ and *ρ*_*k*+1_ be adjacent absolutely super-simple nodes in 𝒢. The *absolutely super-simple structural subnetwork* ℒ′(*ρ*_*k*_, *ρ*_*k*+1_) is the input-output subnetwork consisting of nodes in ℒ (*ρ*_*k*_, *ρ*_*k*+1_) ∪ ℬ, where ℬ consists of all absolutely appendage nodes that are *CS*_*m*_-path equivalent to nodes in ℒ (*ρ*_*k*_, *ρ*_*k*+1_) for some *ι*_*m*_*o*-simple path *S*_*m*_, for some *m* ∈ {1, …, *n*}. That is, ℬ consists of all 𝒜_𝒢_-path components ℬ_*i*_ that are *CS*_*m*_-path equivalent to nodes in ℒ (*ρ*_*k*_, *ρ*_*k*+1_) for some *S*_*m*_, for some *m* ∈ {1, …, *n*}. Arrows of ℒ′(*ρ*_*k*_, *ρ*_*k*+1_) are arrows of 𝒢 that connect nodes in ℒ′(*ρ*_*k*_, *ρ*_*k*+1_). Note that *ρ*_*k*_ is the input node and that *ρ*_*k*+1_ is the output node of ℒ′(*ρ*_*k*_, *ρ*_*k*+1_).

We shall employ the characterisation of super-simple structural subnetworks with respect to each of the input nodes. This was the strategy employed in [38], and hence we will be able to apply directly the results contained in [38, Subsection 3.4.2] to the case of networks with multiple inputs.

First, for *ρ*_*k*_ and *ρ*_*k*+1_ adjacent *ι*_*m*_-super-simple nodes in the core subnetwork 𝒢_*m*_, define as in [38] the *ι*_*m*_*-super-simple structural subnetwork* 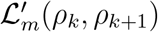 as the input-output subnetwork consisting of nodes in ℒ_*m*_(*ρ*_*k*_, *ρ*_*k*+1_) ∪ ℬ_*m*_, where ℬ_*m*_ consists of all *ι*_*m*_-appendage nodes that are *C*_*m*_*S*_*m*_-path equivalent to nodes in ℒ_*m*_(*ρ*_*k*_, *ρ*_*k*+1_) for some *ι*_*m*_*o*-simple path *S*_*m*_. As usual, arrows of 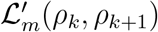 are arrows of 𝒢_*m*_ that connect nodes in 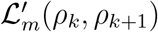.

We notice that [38, Lemma 3.21] is still valid in the current context. Hence, we obtain the following.

##### Lemma 3.7.

*Let ρ*_*k*_ *> ρ*_*k*+1_ *be two adjacent absolutely super-simple nodes. Then* 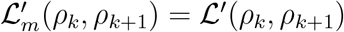, *for every m* = 1, …, *n*.

##### Theorem 3.8.

*Let* 𝒦_*B*_ *be a subnetwork of* 𝒢 *associated with a pleiotropic-structural block B. Then* G *has adjacent absolutely super-simple nodes ρ*_*k*_ *and ρ*_*k*+1_ *such that* 𝒦_*B*_ = ℒ′(*ρ*_*k*_, *ρ*_*k*+1_).

*Proof*. This is a consequence of [38, Thm 3.22]. If *B* is an irreducible pleiotropic-structural block, then it is a structural block associated to each specialized subnetwork 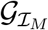. Fix an input ℐ_*M*_. By [38, Thm 3.22], [25, Thm 6.11] and Lemma 3.7, this implies that there exist *ι*_*m*_-absolutely super-simple nodes 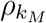 and 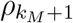 such that 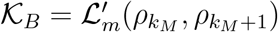 for all *m* such that *ι*_*m*_ is an input node of the specialized subnetwork 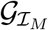. Now, as the input and output nodes of all these networks must be the same, we conclude that there exist absolutely super-simple nodes *ρ*_*k*_, *ρ*_*k*+1_ such that for all *m* = 1, …, *n*, 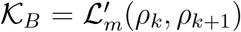. By Lemma 3.7, this means that 𝒦_*B*_ = ℒ′(*ρ*_*k*_, *ρ*_*k*+1_).

Our argument suggests that, as in the case of core networks with only one input [25, 38], the networks supports pleiotropic-structural homeostasis whenever there are more than one absolutely super-simple node.

##### Theorem 3.9.

*If* 𝒢 *has absolutely super-simple nodes other than the output node, then the homeostasis matrix of each absolutely super-simple structural subnetwork corresponds to an irreducible pleiotropic-structural block*.

*Proof*. Consider the adjacent absolutely super-simple nodes *ρ*_*k*_, *ρ*_*k*+1_ in 𝒢. By Lemma 3.5, for every *m* = 1, …, *n*, we have 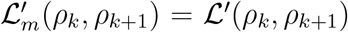. As proved in [38, Cor 3.23], this means that the homeostasis matrix of ℒ′(*ρ*_*k*_, *ρ*_*k*+1_) is an irreducible structural homeostasis block of each subnetwork 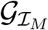. Therefore the homeostasis matrix of ℒ′(*ρ*_*k*_, *ρ*_*k*+1_) is an irreducible pleiotropic-structural homeostasis block.

Finally, we state below a sufficient condition for a core multiparameter network to support coincidental homeostasis. The network of Figure 3(b) shows that condition given below is sufficient, but not necessary, for a core multiparameter network to support coincidental homeostasis.

##### Proposition 3.10.

*Let* 𝒢 *be a multiparameter core network. If none of the input nodes of* 𝒢 *is an absolutely super-simple node, then* 𝒢 *supports at least one coincidental homeostasis type*.

*Proof*. Note that an input node is an absolutely super-simple node if and only if it is the first absolutely super-simple node of the network. Hence, the hypothesis that none of the input nodes of 𝒢 is an absolutely super-simple node is equivalent to: the first absolutely super-simple node of𝒢 is not an input node. First, suppose that pleiotropic homeostasis does not occurs in 𝒢. Then by Proposition 2.8 it follows that coincidental homeostasis must occur in 𝒢. Now suppose that pleiotropic homeostasis does occur in 𝒢. Clearly, as the nodes involved are not absolutely appendage, by Theorem 3.3, they do not belong to a pleiotropic-appendage subnetwork. In addition, none of the input nodes belong to any absolutely super-simple structural subnetwork, as the input nodes are not absolutely super-simple and cannot be between two absolutely super-simple nodes (this is a consequence of [38, Lem 3.15]). Hence, by Theorem 3.8, none of the input nodes belong to a pleiotropic-structural subnetwork. Therefore, each input node belong to a counterweight subnetwork and thus coincidental homeostasis must occur. □

## Acknowledgements

We thank Martin Golubitsky, Ian Stewart, Yangyang Wang, Zhengyuan Huang, Will Duncan, Pedro P. A. C. Andrade and Misaki Yamada for helpful conversations. The research of JLOM and FA was supported by Fundação de Amparo à Pesquisa do Estado de São Paulo (FAPESP) grant 2019/12247-7.

